# EndoC-βH1 multi-genomic profiling defines gene regulatory programs governing human pancreatic β cell identity and function

**DOI:** 10.1101/399139

**Authors:** Nathan Lawlor, Eladio J. Márquez, Peter Orchard, Narisu Narisu, Muhammad Saad Shamim, Asa Thibodeau, Arushi Varshney, Romy Kursawe, Michael R. Erdos, Matt Kanke, Huiya Gu, Evgenia Pak, Amalia Dutra, Sheikh Russell, Xingwang Li, Emaly Piecuch, Oscar Luo, Peter S. Chines, Christian Fuchbserger, NIH Intramural Sequencing Center, Praveen Sethupathy, Aviva Presser Aiden, Yijun Ruan, Erez Lieberman Aiden, Francis S. Collins, Duygu Ucar, Stephen C.J. Parker, Michael L. Stitzel

## Abstract

EndoC-βH1 is emerging as a critical human beta cell model to study the genetic and environmental etiologies of beta cell function, especially in the context of diabetes. Comprehensive knowledge of its molecular landscape is lacking yet required to fully take advantage of this model. Here, we report extensive chromosomal (spectral karyotyping), genetic (genotyping), epigenetic (ChIP-seq, ATAC-seq), chromatin interaction (Hi-C, Pol2 ChIA-PET), and transcriptomic (RNA-seq, miRNA-seq) maps of this cell model. Integrated analyses of these maps define known (e.g., *PDX1, ISL1*) and putative (e.g., *PCSK1, mir-375*) beta cell-specific chromatin interactions and transcriptional *cis*-regulatory networks, and identify allelic effects on *cis*-regulatory element use and expression.

Importantly, comparative analyses with maps generated in primary human islets/beta cells indicate substantial preservation of chromatin looping, but also highlight chromosomal heterogeneity and fetal genomic signatures in EndoC-βH1. Together, these maps, and an interactive web application we have created for their exploration, provide important tools for the broad community in the design and success of experiments to probe and manipulate the genetic programs governing beta cell identity and (dys)function in diabetes.

## INTRODUCTION

Type 2 diabetes (T2D) is a complex disease characterized by elevated blood glucose levels. Ultimately, T2D results when the pancreatic islets are unable to produce and secrete enough insulin to compensate for insulin resistance in peripheral tissues of the body. Individual genetic variation combined with dietary and environmental stressors contribute to disease risk and pathogenesis (Elbein et al., 2012; Franks, 2011; Fuchsberger et al., 2016; Lawlor et al., 2017a; Mohlke and Boehnke, 2015). Genome-wide association studies have identified hundreds of genetic loci associated with T2D and related traits, but extensive work remains to identify the causal/functional variant(s), define their target gene(s), and determine the role(s) of these genes in beta cell identity and function. Several studies have employed (epi)genomic and transcriptomic profiling of human islets (van de Bunt et al., 2015; Dayeh et al., 2014; Fadista et al., 2014; Parker et al., 2013; Varshney et al., 2017; Volkmar et al., 2012), purified/isolated beta cells (Ackermann et al., 2016; Blodgett et al., 2015; Bramswig et al., 2013; Dorrell et al., 2011; Nica et al., 2013), and single cell populations (Baron et al., 2016; Dorajoo et al., 2017; Lawlor et al., 2016; Li et al., 2016; Muraro et al., 2016; Segerstolpe et al., 2016; Wang et al., 2016; Xin et al., 2016) to identify changes in transcriptional regulation and gene expression associated with beta cell-specific functions in individuals with T2D. However, the molecular and physiologic consequences of these alterations and their causal link to beta cell failure and T2D pathogenesis remain largely undefined.

With the recent creation of an immortalized human beta cell line, EndoC-βH1 (Ravassard et al., 2011), islet researchers now possess a necessary tool to experimentally interrogate the molecular mechanisms that govern human beta cell identity and (dys)function in the context of T2D. Since the initial report of their creation, studies utilizing EndoC-ßH1 to build insights into human beta cell regulation and function have grown steadily. These studies have demonstrated that the physiology (e.g., response to glucose, insulin secretion) of EndoC-βH1 cells resembles that of their primary islet counterparts (Andersson et al., 2015; Gurgul-Convey et al., 2015, 2016; Krizhanovskii et al., 2017; Oleson et al., 2015; Teraoku and Lenzen, 2017; Tsonkova et al., 2018) and researchers have used them to identify novel genes involved in human insulin secretion (Ndiaye et al., 2017). In order to motivate further functional studies of human beta cell molecular biology and guide the development of cellular models (e.g., for small molecule screening (Tsonkova et al., 2018)), extensive characterization of the EndoC-βH1 molecular landscape is needed. Here, we completed multi-omic profiling of EndoC-ßH1 cells to extensively map the (1) chromosomal (spectral karyotyping), (2) 3-D epigenomic/chromatin looping (Hi-C (Belton et al., 2012), ChIA-PET (Li et al., 2014)), (3) histone modification (ChIP-seq), (4) chromatin accessibility (ATAC-seq) (Buenrostro et al., 2013), (5) genetic (dense genotyping and imputation), and (6) transcriptomic (RNA-seq, miRNA-seq) signatures of EndoC-ßH1. With these high-resolution maps, we sought to (1) examine gene regulatory programs central to human beta cell identity and function, (2) nominate putative functional variants, putative molecular mechanisms, and target genes underlying T2D, glucose, and insulin genetic associations, and (3) build a publicly available web application for interactive and easy exploration. By comparing these multiomic profiles to those we generated from islets in this study (Hi-C) and in previous studies (Khetan et al., 2017), we identified shared and unique *cis*- regulatory elements (*cis*-RE) and gene expression features. Taken together, these data, the insights gleaned from their analysis, and the research support provided by the web application serve as a high-content resource to enable and guide future functional assessment and molecular studies of beta cell (dys)function.

## Results

### Chromosomal and genetic heterogeneity of EndoC-ßH1

To pursue a precise, comprehensive understanding of the regulatory networks that govern EndoC-βH1/islet beta cell identity and function, we first defined the chromosomal complement and stability of this cell line using spectral karyotyping (SKY) (Figure 1A). SKY analysis of fourteen EndoC-ßH1 metaphase spreads revealed that the number of chromosomes was pseudodiploid (n=46 to 48) (Figure S1). Nearly all metaphases (n = 13/14) had a normal XY sex complement, with only one having a missing Y chromosome, (metaphase 2; Table S1).

**Figure 1.**
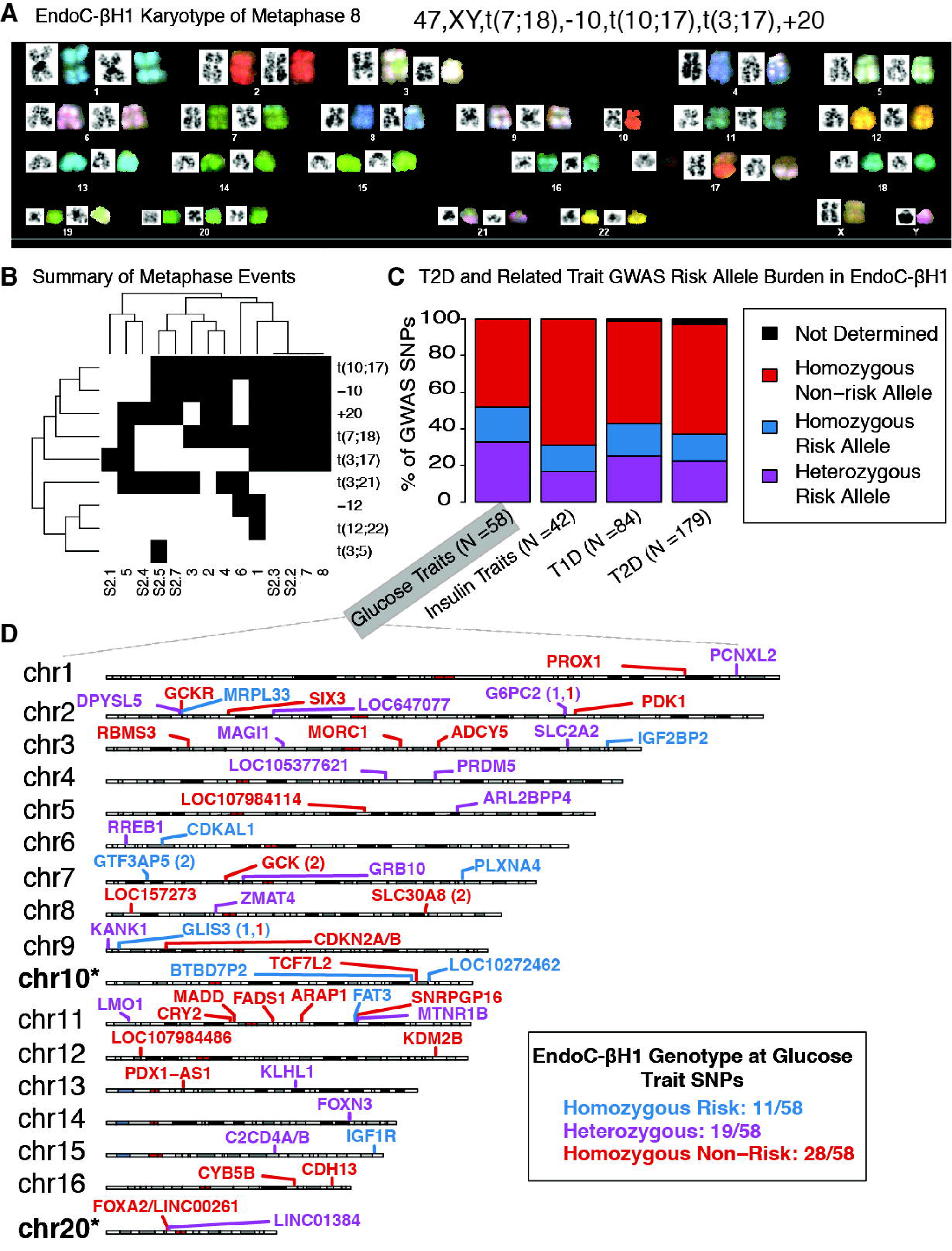
Extensive karyotyping and genotyping of EndoC-ßH1. **(A)** Spectral Karyotyping (SKY) of EndoC-ßH1 for a representative metaphase. **(B)** Summary of the frequency of chromosomal abnormalities across 14 metaphases. Black boxes indicate the presence of an event, while white boxes indicate an absence. **(C)** Bar plots highlighting the risk allele burden of NHGRI-EBI GWAS Catalog diabetes-associated GWAS loci in EndoC-ßH1. T1D = type 1 diabetes, T2D = type 2 diabetes. Glucose traits include fasting plasma glucose and fasting glucose related traits interacting with body mass index (BMI) from the NHGRI-EBI GWAS catalog (MacArthur et al., 2017). Insulin traits include proinsulin and fasting insulin traits interacting with BMI. **(D)** Chromosome cartoons illustrating EndoC-ßH1 genotypes and the reported genes at glucose trait GWAS SNPs. Cases in which independent association signals mapped to the same locus are indicated by the locus name followed by parentheses containing numbers of SNPs with each risk genotype. Chromosomes 10 and 20 are marked with asterisks “*” to indicate that the previously observed copy number alterations (illustrated in Figure 1B) may obfuscate interpretation of variant genotypes on these chromosomes.

The most common autosomal aberrations in EndoC-ßH1 included chromosome 20 gains (n=11/14 metaphases) and chromosome 10 losses (n=10/14). Both of these were also detected as copy number changes by comparative genomic hybridization (CGH) analysis of the cell line (Univercell-Biosolutions). As summarized in Figure 1B and Table S1, we also noted recurrent 10;17 (11/14 metaphases), 7;18 (10/14 metaphases), 3;17 (7/14 metaphases), and 3;21 (7/14 metaphases) chromosomal translocations as well as rarer 12;22 (metaphase 1) and 3;5 (metaphase S2.5) translocations and loss of chromosome 12 (2/14 metaphases; Table S1). Together, these results emphasize that although EndoC-ßH1 is largely diploid, vigilance and caution is warranted when completing and interpreting experiments involving genes or *cis*- REs on chromosomes 3, 7, 10, 17, 18, 20, and 21.

### Delineation of T2D- and related metabolic trait-associated GWAS SNP genotypes in EndoC-ßH1

Genome wide association studies (GWAS) have identified hundreds of index and linked single nucleotide polymorphisms (SNPs) representing putative causal variant(s) at hundreds of loci (Mahajan et al., 2018) associated with genetic risk of T2D and changes in associated quantitative traits (e.g., fasting glucose, insulin, and proinsulin levels). We completed dense genotyping and imputation of EndoC-ßH1 (Materials and Methods) to determine the genotypes at approximately 2.5 million sites genome-wide (MAF > 1%), including disease-associated SNPs. First, we overlapped EndoC-ßH1 genotypes with NHGRI/EBI GWAS cataloged (MacArthur et al., 2017) (Materials and Methods) single lead SNPs associated with glucose levels (fasting glucose), insulin levels (fasting insulin, proinsulin levels), type 1 diabetes (T1D), or T2D. EndoC-ßH1 exhibited homozygous non-risk genotypes at >50% of these SNPs (Figure 1C; Table S2). For approximately 20% of analyzed GWAS loci, EndoC-ßH1 possessed a heterozygous genotype, including rs10830963 at the *MTNR1B* locus (chr11) and rs7173964 at the *C2CD4A/4B* locus (chr15) (Figure 1D; Table S2). Overlap with T2D-associated SNPs (n=6,725 total, constituting 403 unique signals) reported in the most recent meta-analysis (Mahajan et al., 2018) revealed a similar genotype distribution, in which EndoC-ßH1 was heterozygous for approximately 30% (n=119/403; Table S2) of T2D signals. These unique signals represent attractive candidates for epigenomic editing to experimentally determine T2D-associated allelic effects on *cis*-RE use in human beta cells.

### EndoC-βH1 epigenome and transcriptome largely resemble those of primary islets but retain fetal/progenitor islet cell signatures

To identify the genome-wide location of EndoC-ßH1 *cis*-REs (Figure 2A), we generated chromatin accessibility maps using ATAC-seq and defined chromatin states (ChromHMM) by completing and integrating ChIP-seq profiles for multiple histone modifications. ATAC-seq identified 127,894 open chromatin sites in EndoC-ßH1. Qualitative comparison of EndoC-βH1 open chromatin and chromatin state maps to those in primary human islets (Khetan et al., 2017; Varshney et al., 2017) revealed that the genomic architecture for well-known islet-specific loci such as *PCSK1* (Figure 2A), *PDX1*, and *NKX6-1*, was remarkably similar in both, suggesting EndoC-βH1 cells effectively recapitulate beta cell *cis*-regulatory landscapes.

**Figure 2.**
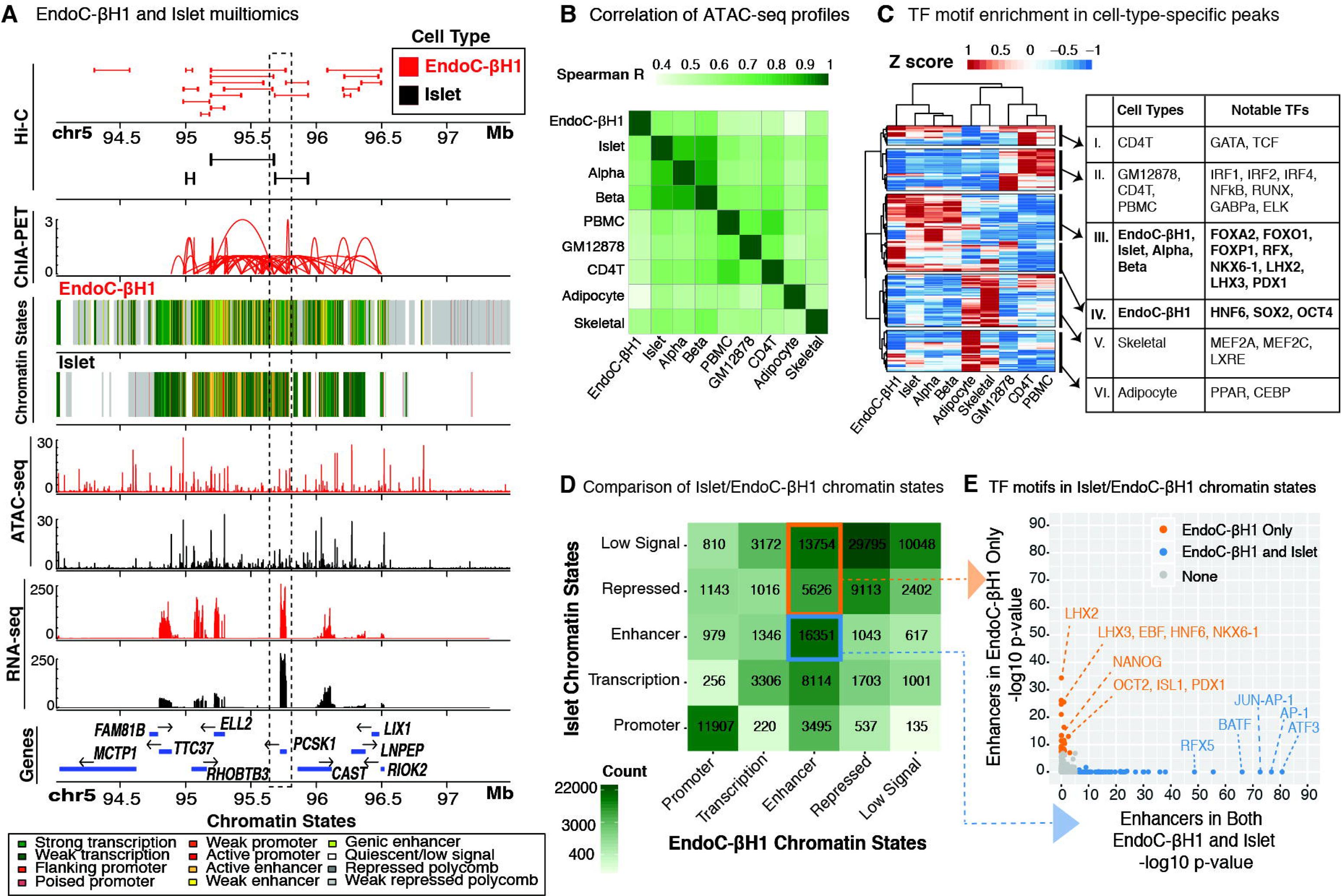
Multi-omic comparative analysis of EndoC-ßH1 and human pancreatic islets. **(A)** Integrated view of the EndoC-ßH1 and human islet (epi)genomic and transcriptomic features surrounding the *PCSK1* locus on chromosome 5. Histone modification ChIP-seq data from EndoC-ßH1, human islets, and 5 Epigenome Roadmap cell types/tissues (Roadmap Epigenomics Consortium et al., 2015) were jointly analyzed to determine ChromHMM-based chromatin states in a uniform manner. **(B)** Spearman correlation between EndoC-ßH1 ATAC-seq profiles and their corresponding profiles from islets, sorted alpha or beta cells, and other cell types and tissues (Methods). PBMC=peripheral blood mononuclear cells, GM12878 = B-lymphoblast cell line, CD4T = CD4^+^ T immune cell, skeletal = skeletal muscle, Alpha = primary islet alpha cells, Beta = primary islet beta cells. EndoC-ßH1 exhibits greatest similarity to islets and their cellular constituents. **(C)** Heat map illustrating z-scores of HOMER enrichment p-values for transcription factor motifs in cell-type-specific OCRs. **(D)** Comparison of chromatin states between EndoC-ßH1 and human islets. Blue box highlights putative enhancer *cis*-REs in both EndoC-ßH1 and human islets; orange box indicates putative EndoC-ßH1 enhancers that are repressed in islets. **(E)** Transcription factor (TF) motifs enriched in genomic regions containing putative enhancer *cis*-REs in both EndoC-ßH1 and islets (blue) or EndoC-ßH1 only (orange). Points in grey denote TFs that are not enriched in either category.

We further compared each ATAC-seq dataset to those from primary islets (Khetan et al., 2017; Varshney et al., 2017), sorted beta/alpha cells (Ackermann et al., 2016), and other primary cell types including adipocyte (Khetan et al., 2017), skeletal muscle (Scott et al., 2016), peripheral blood mononuclear cells (PBMC) (Ucar et al., 2017), and CD4+ T cells (Buenrostro et al., 2013). We identified a total of 269,701 open chromatin regions (OCRs) across all cell types analyzed (Materials and Methods). Among all studied cell types, EndoC-βH1 ATAC-seq profiles were most similar to those of beta cells (Figure 2B, Spearman R = 0.67) and islets (Figure 2B, Spearman R = 0.64).

Next, we compared OCRs and chromatin states to determine where and to what extent EndoC-ßH1 chromatin states recapitulated those of human islets, their constituent cell types, or other metabolic tissues. EndoC-ßH1, islet, and beta cell OCRs were commonly enriched for binding sites of transcription factors (TFs) implicated in islet cellular identity and function (Figure 2C; Group III: FOXA2, FOXO1, RFX, NKX6-1, PDX1). EndoC-ßH1 OCRs also showed exclusive enrichment of sequence motifs that correspond to TFs that regulate pluripotency and pancreatic progenitor states (Figure 2C; Group IV: HNF6, SOX2, OCT4), likely reflecting the fetal origin/derivation of EndoC-ßH1. At EndoC-ßH1 ATAC-seq OCRs, promoter annotations (from ChromHMM; Materials and Methods) were widely conserved across EndoC-ßH1 and other cell types including islets as expected (Figure S2A; centered Pearson correlation > 0.95; Materials and Methods). In contrast, enhancers, which often encode cell-specific transcriptional regulatory elements (Heinz et al., 2015), at EndoC-ßH1 ATAC-seq OCRs were most comparable between islet and EndoC-ßH1 (Figure S2A; centered Pearson correlation ∼ 0.71; black point).

To further assess similarities and differences between islet and EndoC-ßH1 epigenomes, we investigated the proportions and features of chromatin states that were preserved or disparate between them. Unsurprisingly, a large proportion of promoters (11,907/19,482; ∼61%) were preserved (Figure 2D) which included motifs for a variety of TFs from the ETS family (e.g., ELK4, ETS, ELF1; Table S3) with established roles in cellular differentiation, proliferation, and apoptosis (Findlay et al., 2013). Regions annotated as repressed in both islets and EndoC-ßH1 were enriched for CTCF and BORIS binding motifs (Table S3), DNA binding proteins known to bind and establish transcriptional insulators at chromatin territory boundaries. 16,351/51,325 putative enhancers (defined via ChromHMM) were shared between islets and EndoC-ßH1 (Figure 2D; blue box) and showed strong enrichment for general (ATF3, AP-1, JUN) TFs (Figure 2E, blue dots; Table S3) relative to all enhancer regions. Interestingly, we observed a substantial number of EndoC-ßH1 enhancers that were annotated as quiescent/repressed in islets (n = 19,380) (Figure 2D; orange box). Relative to all enhancers, these sites were enriched for TFs controlling pluripotency (OCT2, NANOG) (Sokolik et al., 2015; Tantin, 2013), pancreatic development/lineage specification (HNF6, ISL1) (Zhang et al., 2009), and beta cell fate determination (PDX1, NKX6-1) (Thompson and Bhushan) (Figure 2E, orange dots; Table S3). Based on these findings, it is possible that these regions represent fetal or developmental *cis*-REs that are active in the fetal-derived EndoC-ßH1 and inactive in adult islets composed of mature beta cells. Nonetheless, a significant number of *cis*-REs (n = 16,351) are conserved between EndoC-ßH1 and human islets.

Next, we measured EndoC-ßH1 gene expression using RNA-seq and compared it to RNA-seq profiles of islets and other cell types/tissues (Figure S2B). As anticipated, the EndoC-ßH1 transcriptome most strongly correlated with transcriptomes of islets (R = 0.87) and primary beta cells (R=0.86) among all tissues/cells tested. Of the 27,564 protein coding/lincRNA genes considered, 11,554 were expressed in EndoC-ßH1 and 12,231 genes were expressed in islet (with 10,473 genes expressed in both). Similarly, EndoC-βH1 small non-coding RNA (miRNA) profiles resembled human islets more than the other tissues (adipose, skeletal muscle; Figure S2C) in a principal component analysis (PCA). In particular, PC1 loadings were highly correlated with key islet miRNAs including, miR-375 (Table 1), a critical regulator of beta cell mass and identity (Eliasson, 2017), whereas PC2 stratified primary tissue (adipose, skeletal muscle, islet) from immortalized cells (EndoC-βH1). Consistent with the PCA, miRNA expression levels in EndoC-βH1 and islets were highly correlated (R = 0.779; Figure S2D), and the vast majority of the most highly expressed miRNAs in EndoC-βH1 have been reported previously to be enriched in primary human beta cells relative to whole islets (Table 1) (Bunt et al., 2013). Together, these chromatin accessibility, chromatin state, gene expression, and small RNA expression analyses reveal substantial conservation between the transcriptional regulatory and gene expression landscapes of EndoC-ßH1 and primary islets.

### Hi-C profiling of EndoC-ßH1 and human islets reveals beta cell-specific chromatin looping domains

Next, we sought to determine spatial chromatin organization and identify chromatin domains in EndoC-ßH1 and islets using Hi-C. We generated Hi-C maps for EndoC-ßH1 and islet cells with approximately 6 billion reads each. The maps have 1.9 billion contacts and 1.5 billion contacts, respectively. Using Juicer (Durand et al., 2016a) (Material and Methods), we identified 9,100, 2,580, and 9,448 Hi-C loops in EndoC-ßH1, human islets, and GM12878 (human lymphoblastoid cell line) (Rao et al., 2014), respectively. Together, this represents 19,428 independent DNA-DNA loops. Aggregate Peak Analyses (APA) (Rao et al., 2014) (Figure 3A, top plots) revealed that chromatin looping sites (anchors) were comparable in EndoC-ßH1, islets, and GM12878 for the majority of (> 90%) the total chromatin loops (n=19,428/21,128). Consistent with previous studies (Rao et al., 2014; Vietri Rudan et al., 2015), CTCF and CTCFL/BORIS DNA binding motifs were overwhelmingly enriched (p<1e-229, p<1e-114, respectively) among all Hi-C anchor sequences (Table S4), verifying that general 3-D chromatin structures and loops are preserved between different mammalian tissues and cell types. Importantly, however, we detected 1,078 chromatin loops that were exclusively present in EndoC-ßH1 compared to GM12878 (Figure 3A, bottom plots), a subset of which (n=50) was independently detected in human islet Hi-C data.

**Figure 3.**
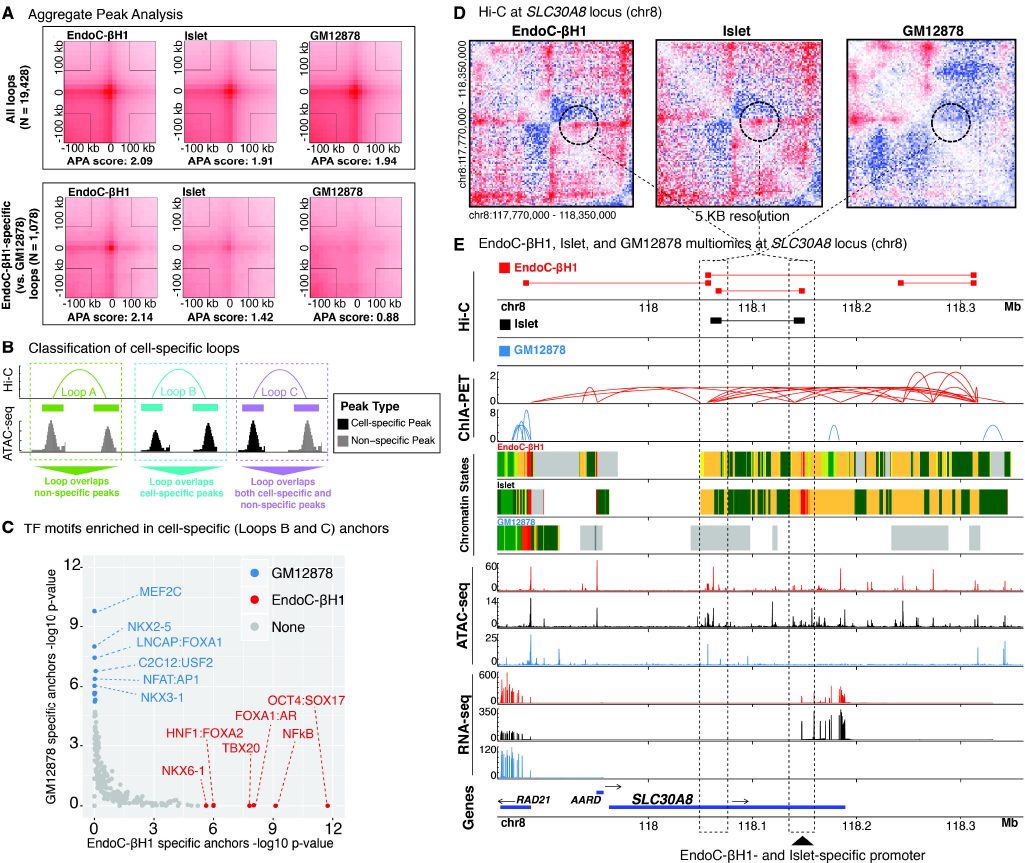
Generating a genome-wide map of looping in EndoC-ßH1 and human pancreatic islets (Hi-C) **(A)** Aggregate peak analysis (APA) plots showing the total signal across all loops (top three panels) and EndoC-ßH1-specific loops (bottom three panels) in EndoC-ßH1 (left), human islet (center), and GM12878 (right) cells. Of note, islets exhibit similar contact point enrichments at EndoC-ßH1-specific peaks compared to GM12878. **(B)** Cartoon illustrating the different classes of Hi-C loops between common (gray peaks) or cell-specific (black peaks) ATAC-seq OCRs. **(C)** Transcription factor motifs enriched in GM12878 (blue) or EndoC-ßH1 (red) Hi-C looping anchors that overlap cell-specific ATAC-seq peaks (Loop classes B and C in panel B above). **(D)** Hi-C contact maps highlighting a specific loop at the *SLC30A8* locus (denoted by dotted black circle) observed in both EndoC-ßH1 (left) and primary human islets (center) but absent in GM12878 (right). **(E)** Multi-omics view of Hi-C, ChIA-PET (Pol2), chromatin states, ATAC-seq, RNA-seq, and gene tracks at the *SLC30A8* neighborhood containing the Hi-C contact point highlighted in panel D. Tracks corresponding to EndoC-ßH1, human islet, and GM12878 are colored red, black, and blue, respectively.

To further study cell-specific loops, we subdivided EndoC-ßH1 and GM12878 differential Hi-C loops into three categories based on the cell type-specificity of the ATAC-seq OCRs they bring into physical proximity (Figure 3B): (A) loops between two non-specific OCRs, (B) loops between two cell-specific OCRs, or (C) loops between one cell-specific OCR and one non-specific OCR. Class B/C loops were classified as cell-specific and further studied. Comparison of EndoC-ßH1-specific (n = 315) and GM12878-specific (n = 308) loops revealed a strong bias for cell-specific TF binding at anchors (Figure 3C). In EndoC-ßH1-specific anchors, we observed enrichment for TFs involved in beta cell differentiation and function (NKX6-1, FOXA2, FOXA1) (Thompson and Bhushan) as well as OCT4, a key regulator for early embryo development (Le Bin et al., 2014; Wu and Schöler, 2014), while GM12878-specific anchors were enriched for TFs necessary for B cell proliferation and activation (MEF2C, NFAT) (Herglotz et al., 2016; Peng et al., 2001). Furthermore, genes adjacent to EndoC-ßH1-specific anchors (Materials and Methods) were most enriched (Hypergeometric FDR-adjusted p-value < 0.05) for islet-associated GO terms including insulin secretion, glucose homeostasis, and neuronal/endocrine development (Figure S3A; Table S4 for complete results). For several genes affiliated with these GO terms, such as *SLC30A8*, which encodes a zinc efflux transporter closely involved with zinc ion sequestering and ultimately insulin secretion (Mitchell et al., 2016), we observed striking similarities in Hi-C contact frequencies between islet and EndoC-ßH1 (Figure 3D). In contrast, we observed far fewer chromatin loops and large spans of polycomb-repressed and/or quiescent chromatin for these loci in GM12878 cells (Figure 3E).

Approximately 50% (4,543/9,100) of EndoC-ßH1 loop anchors overlapped EndoC-ßH1 ATAC-seq OCRs, of which 44% (n=1,987/4,543) of these occurred between promoter and enhancer elements (Figure S3B). Of these loops, we observed a substantially higher proportion (64%; 376/587) of EndoC-ßH1-specific loops that overlapped EndoC-ßH1 stretch enhancers (Fisher’s exact test p-value < 4.23 e-42) compared to that of non-specific loops (34%; 1,358/3,956). To examine the functional specificity of these loops, we overlapped chromatin state (ChromHMM) information from EndoC-ßH1 and 27 other tissue/cell types for all EndoC-ßH1 Hi-C. For each cell type, we determined the percent of Hi-C anchors that contained the same chromatin state as EndoC-ßH1 (Materials and Methods). Islets had the highest percent of identical chromatin states to EndoC-ßH1 at Hi-C anchor sites among all tested tissues/cell types, especially promoter-enhancer loops (Figure S3C; orange line plot). These findings enumerate regions of cell-specific chromatin looping associated with islet development and function and indicate that EndoC-ßH1 forms cell type-specific chromatin domains/territories highly similar to primary human islets.

### EndoC-ßH1 Pol2 ChIA-PET identifies beta cell *cis*-regulatory hubs

To map functional *cis*-regulatory networks, we completed RNA polymerase (Pol 2) ChIA-PET (Li et al., 2017b) in EndoC-ßH1 to identify 25,336 putative Pol2-mediated chromatin interactions. We further filtered these interactions, retaining only those for which both interacting sites (ChIA-PET anchors) overlapped EndoC-ßH1 ATAC-seq OCRs, resulting in 16,756 putative *cis*-regulatory interactions (Materials and Methods). As shown in Figure 4A, the overwhelming majority of Pol2-mediated chromatin interactions linked active enhancer and active promoter chromatin states to themselves and each other. Importantly, ChIA-PET detected EndoC-ßH1-specific loops (Figure 4B, compare EndoC-ßH1, GM12878, and K562 ChIA-PET tracks) coinciding with those previously reported in targeted 4C-seq analyses of human islets (Pasquali et al., 2014), including the *ISL1* (Figure 4B; n = 8 sites denoted by asterisks, and *PDX1* (Figure S4A, n = 9 sites) loci.

**Figure 4.**
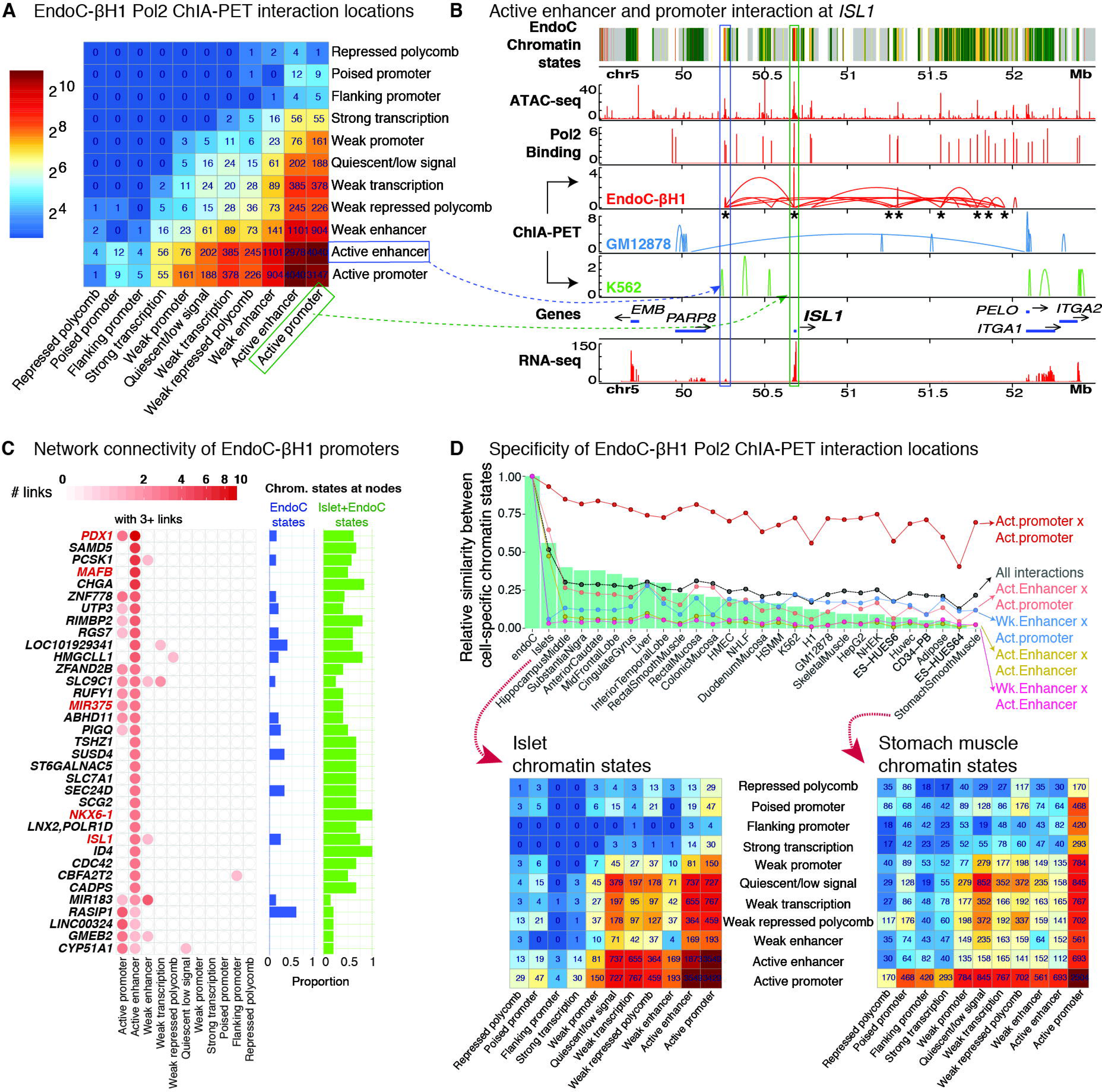
RNA Polymerase 2 ChIA-PET identifies chromatin interactions in EndoC-ßH1. **(A)** Heatmap showing the chromatin states of EndoC-ßH1 ChIA-PET interaction nodes. **(B)** Example of a Pol2 ChIA-PET interaction between active enhancer (blue box) and active promoter (green box) *cis*-REs in the *ISL1* locus on chromosome 5. Asterisks under EndoC-ßH1 ChIA-PET interactions (red) indicate interacting sites in the *ISL1* locus detected in human islet 4C-seq analyses (Pasquali et al., 2014). **(C)** ChIA-PET network connectivity of gene promoters in EndoC-ßH1 containing at least 3 interactions with other regulatory elements. For each gene, the number of connections between other regulatory elements (e.g. active enhancer, weak enhancer) and the proportion of links in which the chromatin states are EndoC-ßH1-specific (blue) or identical in both human islet and EndoC-ßH1 (green) are shown in bar plots on the right. **(D)** (Top) Bar plot illustrating the proportions of chromatin states at the Pol2 ChIA-PET interacting sites (nodes) shared between EndoC-ßH1, islets, and additional Epigenomics Roadmap tissues and cell lines. (Bottom) Heatmap demonstrating the chromatin states of EndoC-ßH1 Pol2 ChIA-PET interacting sites (nodes) in islets (left) or stomach smooth muscle (right).

In addition to replicating interactions previously detected by 4C-seq in human islets, Pol2 ChIA-PET identified genome-wide interactions that encapsulate hundreds of promoter-promoter and putative promoter-enhancer interactions (Figure 4C; Materials and Methods). These include extensive Pol 2 interactions in loci containing genes crucial for beta cell identity and development such as *PDX1, ISL1, NKX6-1, MAFB*, and *miR375* (Figure 4C, red text). As shown in Figures 4C and S4A, multiple interactions were detected between the *PDX1* promoter and classically described essential *PDX1* transcriptional enhancer sequences (n=7/14 enhancer interactions), which contain binding sites for islet TFs such as FOXA2 (Gao et al., 2008; Gerrish et al., 2004). During embryonic development, C57BL/6 mouse pancreata displayed a transition in expression of MafB to MafA (Nishimura et al., 2006) suggesting that these two factors are tightly involved in beta cell differentiation and function. The high degree of connectivity in *MAFB* (vs. that of *MAFA*) may therefore reflect the fetal/naive state of EndoC-ßH1 cells. *miR375*, a small non-coding RNA, possessed multiple connections to active promoter and enhancer elements (Figure 4C; red text), consistent with its role as a post-transcriptional regulator of genes involved in beta cell development/differentiation and insulin secretion/exocytosis (Eliasson, 2017).

Interestingly, *INSM1*, a gene necessary for pancreatic endocrine cell differentiation (Osipovich et al., 2014), harbored the most connections in EndoC-ßH1 (n = 97 total interactions, n = 7 between active promoters and enhancers; Figure S4B). Other genes linked by ChIA-PET interactions are involved in insulin processing and secretion including *PCSK1*, one of the prohormone convertases that catalyzes (pro)insulin processing; *RIMBP2*, whose protein mediates formation of a complex for polarized accumulation and exocytosis of insulin granules (Fan et al., 2017); *RGS7*, a critical regulator of muscarinic-stimulated insulin secretion (Wang et al., 2017); and *CDC42*, which is essential for second phase insulin secretion (Wang et al., 2007). Additionally, genes implicated in the protection and management of stress were highly connected in EndoC-ßH1 ChIA-PET interactions. Notable candidates were *ZFAND2B* whose induction helps protect against human beta amyloid peptide toxicity/accumulation in a *C. elegans* transgenic Alzheimer’s disease model (Hassan et al., 2009); *SUSD4*, a complement inhibitor and tumor suppressor that modules ER stress; and CD59, which is required for mediating exocytosis events facilitating insulin secretion (Blom, 2017; Krus et al., 2014). Finally, *TSHZ1*, a PDX1 target gene whose expression levels were notably lower in human islet donors with T2D (Raum et al., 2015), harbored five links to active enhancer elements in EndoC-ßH1, suggesting that perturbation of the *cis*-regulatory networks identified herein may contribute to T2D pathogenesis.

Finally, we sought to study to what extent the putative *cis*-regulatory networks detected in EndoC-ßH1 may be preserved in islets and other cell types. However, due to a limited availability of ChIA-PET data in human islets and other relevant tissues, we decided to use the chromatin interaction sites determined by EndoC-ßH1 ChIA-PET and compare the functional annotations (ChromHMM state annotations) at these loci across 27 different tissue/cell types. Overall, aggregate counts of these chromatin state interactions for each cell type were most similar between EndoC-ßH1 and islet (Figure 4D; green bar plots; Materials and Methods). ChIA-PET interactions between regions annotated as active promoters in EndoC-ßH1 were similarly annotated as active promoters across the 27 other cell/tissue types (Figure 4D; Act. promoter × Act. promoter red line plot; Materials and Methods). In contrast, the majority of active enhancers interacting in EndoC-ßH1 were marked as active enhancers only in human islets (Figure 4D, Act. Enhancer × Act. Enhancer yellow line plot). Multidimensional scaling of all cell/tissue chromatin state annotations at EndoC-ßH1 ChIA-PET interacting sites also reaffirmed a high similarity between EndoC-ßH1 and islets (Figure S4C) consistent with a strong conservation of active enhancer state annotations as previously observed (Figure 4D; line plots). These results suggest that these interactions may be reflecting beta cell *cis*-regulatory hubs. Indeed, we observed a large proportion of ChIA-PET interactions (6,904/16,756 (41%)) whose anchors overlapped islet stretch enhancers implicating that these interactions may encompass key islet functional chromatin domains.

### Integration of EndoC-ßH1 genotype and 3-D genomic interaction maps identifies allelic imbalance at beta cell-specific *cis-REs*

We and others have demonstrated that genetic variants, including those associated with T2D and other quantitative measures of islet (dys)function, can alter *cis*-RE use (chromatin accessibility quantitative trait loci; caQTL) (Karczewski and Snyder, 2018; Timpson et al., 2018) and target gene expression (expression quantitative trait loci; eQTL (van de Bunt et al., 2015; Fadista et al., 2014; Varshney et al., 2017)). Recently, approaches have been used to assess allelic effects on these molecular features at heterozygous sites within a single sample. We applied these allelic imbalance (AI) analyses in EndoC-ßH1 to assess genetic effects on beta cell *cis*-regulatory networks and gene expression. To identify instances of AI in EndoC-ßH1, we considered approximately 2 million heterozygous SNPs (Materials and Methods) and examined their biases within OCRs (ATAC-seq peaks), active enhancer elements (H3K27ac peaks), or expressed genes (RNA-seq; Figure 5A). Subsequent analyses revealed that <10% of all SNPs occurring in OCRs and enhancer elements showed significant AI (Figure 5A; Part I; FDR < 10%). We observed ∼25% of SNPs with gene expression AI (Figure 5A). When considering variants with adequate coverage in both EndoC-ßH1 ATAC-seq OCRs and H3K27ac-marked enhancer regions (n = 1,734 SNPs), we noted a positive correlation (R = 0.2) in the corresponding AI ratios (Figure S5A) suggesting the potential for coordinated chromatin accessibility and histone modifications at these *cis*-regulatory sites. In total, 119/403 (∼30%) T2D-associated signals overlapped EndoC-ßH1 *cis*-REs (Table S2), of which only 34/119 (∼29%) of these unique signals were heterozygous in EndoC-ßH1 and potentially amenable to allelic analyses. GREGOR (Schmidt et al., 2015) enrichment analysis of these same 403 signals and all corresponding SNPs in high linkage disequilibrium (R^2^ > 0.8), identified significant overlap (p-value < 1 e-7) in EndoC-ßH1 ATAC-seq (n = 67 SNPs) OCRs. These results suggest that EndoC-ßH1 *cis*-REs modestly encompass established T2D signals.

**Figure 5.**
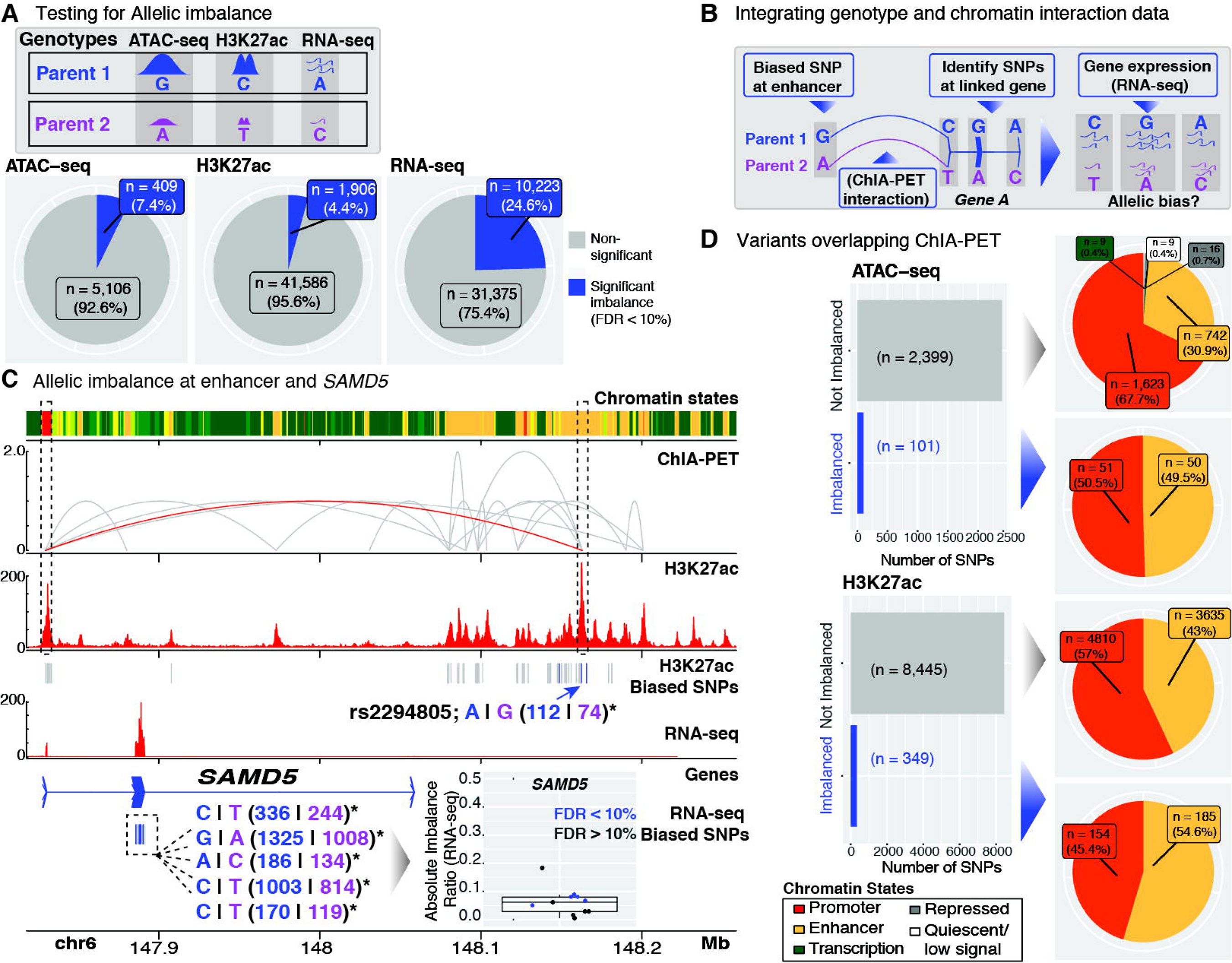
Allelic effects on EndoC-ßH1 transcriptional regulatory features. **(A)** EndoC-ßH1 genotype information was integrated with ATAC-seq, H3K27ac, and RNA-seq data to identify sequence variants altering *cis*-RE accessibility/activity (ATAC-seq, H3K27ac) or mRNA levels (RNA-seq) in EndoC-ßH1. Pie charts summarize the proportions of variants exhibit significant allelic imbalances (blue; FDR < 10%) in each of the corresponding –seq profiles. **(B)** Cartoon representation of approach to identify systematic allelic effects on EndoC-ßH1 *cis*-regulatory networks. **(B)** Multi-omic view highlighting allelic effects on the *SAMD5* locus *cis*-regulatory network in EndoC-ßH1. A variant site exhibiting significant allelic imbalance in H3K27 acetylation (denoted by blue arrow) is linked (red ChIA-PET interaction) to the transcription start site (TSS) of *SAMD5.* Within the *SAMD5* locus, five transcribed SNPs exhibited significant allelic bias in gene expression (RNA-seq) in a direction consistent with the H3K27ac allelic bias. **(D)** (Left) Bar plots summarizing the proportions of variants with ATAC-seq/H3K27ac imbalance (blue bars; FDR < 10%) that overlap ChIA-PET interacting loci. (Right) Pie charts specifying the chromatin state (ChromHMM) annotations of the overlapping variants.

Next, we leveraged information from EndoC-ßH1 ChIA-PET interactions to determine potential allelic effects on *cis*-regulatory networks and target gene expression. To achieve this, we (1) identified SNPs overlapping enhancers with AI (H3K27ac/ATAC-seq), (2) determined if the SNP linked (via ChIA-PET) to the transcription start site (TSS) of a gene, and (3) assessed if promoter or transcribed SNPs in the predicted target gene exhibited allelic imbalance in H3K27ac/ATAC-seq or RNA-seq, respectively (Figure 5B). For example, rs2294805 exhibits allelic imbalance in an EndoC-ßH1 enhancer downstream of *SAMD5* and is linked to this gene’s TSS by a ChIA-PET interaction (Figure 5C). Notably, 5/11 transcribed *SAMD5* SNPs exhibited significant AI in RNA-seq gene expression data. In all cases, one parental chromosome (denoted in blue) was consistently overrepresented in both H3K27ac ChIP-seq and RNA-seq data. Although the exact role of *SAMD5* in human islets has not been described, expression of this gene is high in adult alpha cells, but absent in adult beta cells (Lawlor et al., 2016; Segerstolpe et al., 2016; Wang et al., 2016). *SAMD5* has been recently identified as a marker of peribiliary gland (PBG) cells (Yagai et al., 2017), and PBG stem cells have been documented to differentiate into glucose-responsive pancreatic islets (Cardinale et al., 2011). Our data highlights *SAMD5* as one of the most highly-connected loci in EndoC-ßH1 (Figures 4C and S4B; n = 35 ChIA-PET connections between the gene and active enhancers; n = 5 interactions to other active promoters), and a potential novel *cis*-regulatory hub for fetal beta/islet cell development. Further manipulation of the regulatory networks in this locus may provide greater insight of its putative roles in islet cell differentiation and function.

Overall, 2,500/5,515 (∼45%) and 8,794/43,492 (∼20%) of heterozygous SNPs that passed coverage thresholds (Materials and Methods) in ATAC-seq and H3K27ac *cis*-REs respectively overlapped a ChIA-PET anchor (Figure 5D; bar plots). For both datasets, < 5% of SNPs demonstrated significant AI, which were present in active promoter and enhancer regions of the genome (Figure 5D; pie charts). SNPs with significant H3K27ac and ATAC-seq AI were enriched for active enhancer elements compared to promoter elements (Figure 5D). We also observed 21/50 (42%) and 91/185 (∼49%) of enhancer SNPs (Figure 5D; yellow portion of pie charts marked with blue arrow) with significant ATAC-seq and H3K27ac AI, respectively, linked to target gene promoters via ChIA-PET (Table S5). To determine if the allelic effects on EndoC-ßH1 *cis*-regulatory networks extended to primary islets, we examined the direction-of-effect of these SNPs on steady state expression in human islet eQTLs (Varshney et al., 2017). Islet eQTL Z-scores (Materials and Methods) were most correlated to the H3K27ac allelic ratio at SNPs where the human islet eQTL target gene (i.e., the gene whose expression in human islets is influenced by a genetic variant) matched that of the EndoC-ßH1 ChIA-PET target gene (i.e., the gene linked to the enhancer by EndoC-ßH1 ChIA-PET as depicted in Figure 5B) (n = 42/91) (Figure S5B red points; R = 0.32), as opposed to those genes in the locus that were not linked to the enhancer by ChIA-PET (Figure S5B, grey points; R=0.17).

Importantly, these analyses suggest that integrated EndoC-ßH1 omics analyses provide molecular insights into diabetes genetics (e.g., GWAS). For example, a T2D-associated index SNP rs57235767, for which the ‘C’ risk allele exhibited reduced EndoC-ßH1 H3K27ac counts, exhibited consistent down-regulation of the ChIA-PET-predicted target gene *C11orf54* (Figure 5SB; asterisk) expression in islet cohorts (Fadista et al., 2014; Varshney et al., 2017). Similarly, the T2D risk allele of rs3807136, a SNP in linkage disequilibrium (R^2^ > 0.8) with the index SNP rs2268382, displayed a higher proportion of H3K27ac counts in EndoC-ßH1 and increased expression of the predicted target gene, *CEP41*. These human islet SNP-gene interactions (that are also recapitulated in EndoC-ßH1 cells) represent high priority targets for (epi)genomic modification and should assist efforts to decrypt the factors contributing to T2D pathogenesis.

## Discussion

Here, we report extensive multi-omic mapping and integrated analysis of cytogenetic (karyotyping), large scale chromatin structural conformation (Hi-C), *cis*-regulatory networks (ChIA-PET), histone mark (ChIP-seq), chromatin accessibility (ATAC-seq), genetic (genotyping), and gene expression (RNA-seq) information in EndoC-ßH1 human beta cells. For convenient and interactive browsing of this data, we have created an R shiny web application (Chang et al., 2018) which is publicly available at… These data and the browser application will serve as a resource for future studies to explore the complex beta cell regulatory programs uncovered in this study and to guide targeted studies of regulatory networks, genes, and pathways of interest.

Spectral karyotyping revealed chromosomal heterogeneity among individual cells in the EndoC-ßH1 population. These included copy number variation, such as chromosome 20 gain and chromosome 10 loss, that has been identified previously by array comparative genomic hybridization analyses, as well as previously unappreciated structural alterations, including chromosome 10:17 and 3:21 translocations, that are frequent within the population. Consistent with spectral karyotyping, we observed enhanced contact frequency between chromosomes 3 and 21 in EndoC-ßH1 Hi-C maps (Juicebox (Durand et al., 2016b) link…) suggesting that these two technologies may complement one another to identify cell line abnormalities. As evident by other less prevalent chromosomal aberrations among the population, it is possible that this cell line may continue to evolve with continued passaging. Thus, caution should be taken, and these aberrations should be considered, in future (epi)genome editing or EndoC-ßH1 molecular and functional experiments involving genes or regulatory elements on these chromosomes.

Overall, comparative analyses of omics profiles indicate substantial similarity between EndoC-ßH1, islet, and primary beta cell transcriptomes (Figure 2B; Pearson R > 0.86). EndoC-ßH1 open chromatin profiles were only modestly correlated with islet (R = 0.64) and primary beta (R = 0.67) cells, highlighting potential drift between the cell line and primary islet cells at the epigenetic level. These include approximately 19,000 putative EndoC-ßH1 enhancers that are annotated as quiescent or polycomb repressed in human islets. Our analyses revealed that these sites contain potential binding sites for TFs with important roles in beta cell development and pancreatic precursor fates and functions (e.g., NKX6-1, PDX1, ISL1, HNF6) (Thompson and Bhushan) and pluripotency (e.g., NANOG, OCT2) (Sokolik et al., 2015; Tantin, 2013). These features may reflect the fetal nature of EndoC-ßH1 cells, their transformed nature, or both.

Using Hi-C to map higher order chromatin structure in EndoC-ßH1 and a corresponding map for a representative human islet, we defined islet/beta cell chromatin domains and territories. Consistent with previous findings (Rao et al., 2014), the overall spatial chromatin organization was similar across EndoC-ßH1, islet, and GM12878 cells (Figure 3A; top panel) and all Hi-C anchors were enriched for classic TFs involved in chromatin modeling (e.g., CTCF, BORIS; Table S4). Importantly, however, Hi-C analyses also identified approximately 1,078 islet/beta cell-specific chromatin domains, several of which were evident in both EndoC-ßH1 and primary human islets (Figure 3). These cell-specific chromatin territories were enriched for beta cell-specific TFs (Figure 3C) and brought into close physical proximity genes linked to islet-associated biological process gene ontology terms (Figure S3A) compared to those of GM12878.

Here, we also report the first Pol2 ChIA-PET map of chromatin interactions in EndoC-ßH1, which further refined chromatin territories to reveal functional EndoC-ßH1 *cis*-regulatory networks. In addition to validating chromatin interactions previously reported in 4C-seq analyses at select loci in human islets, Pol2 ChIA-PET identified hundreds of new interactions genome-wide between active promoter and enhancer regions (Figure 4A) potentially involved in the regulation and transcription of dozens of beta cell-specific loci (Figure 4B, C). Comparison of Pol2 ChIA-PET interaction locations in EndoC-ßH1, GM12878, and K562 revealed that the overwhelming majority of interactions at these loci were unique to EndoC-ßH1. Due to high cell input requirements (∼100 million cells) (Li et al., 2017b) for current ChIA-PET library construction protocols, we were unable to validate these findings in human islets. However, consistent with our previous observations between islet and EndoC-ßH1 Hi-C maps (Figure S3C), we noted that chromatin states of ChIA-PET interaction nodes in EndoC-ßH1 were most conserved in islet (Figure 4D, Figure S4C) in comparison to that of 27 other cell/tissue types. Thus, the *cis*-regulatory programs we define for EndoC-ßH1 should provide valuable insight of important transcriptional hubs that drive islet and beta cell identity and function.

Genome-wide association studies have identified hundreds of SNPs that contribute to genetic risk of type 2 diabetes and other quantitative measures of islet dysfunction, such as glucose, insulin, and proinsulin levels (DIAbetes Genetics Replication And Meta-analysis (DIAGRAM) Consortium et al., 2014; Fuchsberger et al., 2016; Mahajan et al., 2018). We and others have linked a subset of SNPs and loci to altered *cis*-regulatory element activity and steady state islet gene expression (van de Bunt et al., 2015; Fadista et al., 2014; Gaulton et al., 2015; Parker et al., 2013; Pasquali et al., 2014; Varshney et al., 2017). For the majority of loci, challenges remain to define the (1) causal or functional SNP, (2) determine its molecular effect on *cis*-regulatory element activity, and (3) identify the putative target gene(s). By combining our densely genotyped/imputed and 3-D chromatin interaction (ChIA-PET) networks in EndoC-ßH1, we sought to identify SNPs with imbalanced expression or *cis*-RE use and link them physically with their target genes. Using this approach, we linked 91/185 (H3K27ac imbalanced) and 21/50 (ATAC-seq imbalanced) SNPs to potential target genes (e.g., rs2294805 -*SAMD5* in Figure 5B). Our ability to assess chromatin interactions between diabetes-associated SNPs and their target genes was modest. Nonetheless, we identified two candidate SNPs (rs57235767, rs3807136) that demonstrated consistent directions-of-effect on epigenetic measures of *cis*-regulatory element activity (e.g., H3K27ac) and target gene expression in EndoC-ßH1 and human islets (Figure S5B) (Varshney et al., 2017). Several factors could underlie the modest frequency of diabetes-associated GWAS SNPs linked by Pol2 ChIA-PET interactions: (1) limited sensitivity of ChIA-PET technology (2) condition/disease specificity of GWAS SNP effects on *cis*-regulatory element use or activity; or (3) condition-specific Pol2 interactions between *cis*-REs and their target genes.

In summation, this study provides the first integrated multi-omic analysis of EndoC-ßH1, a human pancreatic beta cell line with increasing utility and importance to the beta cell and diabetes communities. Integrated analysis of 3-D epigenetic and gene expression information identified chromosomal territories and *cis*-regulatory networks governing beta cell identity and function. Overall, comparison of EndoC-ßH1 (epi)genomic and 3-D chromatin profiles with those of human islets verified common signatures of gene expression, TF binding, and *cis*-RE use. These analyses also highlighted genomic discrepancies between EndoC-ßH1 and their primary cell counterparts, presumably reflecting the fetal/embryonic origin of the cell line and/or its transformed state. Integration of EndoC-ßH1 *cis*-regulatory maps with genome-wide genotype information nominated target genes and identified SNP allelic effects on transcriptional regulatory networks, including a subset of T2D-associated SNPs. Together, the data and tools provided here should serve as helpful guides for comprehensive design of targeted and hypothesis-driven studies of candidate genes, pathways, or *cis*-REs to determine their role(s) in beta cell (dys)function and diabetes.

## Acknowledgements

We thank Jane Cha for aid in graphic design. We thank all members of the Stitzel, Ucar, Parker, Sethupathy, Collins, Ruan, and Lieberman-Aiden labs for helpful discussion and feedback on this study and manuscript. This study was supported by NIH training grant T32 HG00040 (to P.O.), NIH (DP2OD008540, U01HL130010, UM1HG009375), NSF (PHY-1427654), USDA (2017-05741), Welch Foundation (Q-1866), NVIDIA, IBM, Google, Cancer Prevention Research Institute of Texas (R1304), and McNair Medical Institute (to E. L. A.), NIH intramural support from project ZIA-HG000024 to (F.S.C.), NIH grants R00DK092251 (to M.L.S), R00 DK099240 (to S.C.J.P) and support from the American Diabetes Association Pathway to Stop Diabetes Initiator (1-14-INI-07 to S.C.J.P.) and Accelerator (1-16-ACE-to P.S. and 1-18-ACE-15 to M.L.S.) Awards. Opinions, interpretations, conclusions, and recommendations are solely the responsibility of the authors and do not necessarily represent the official views of the National Institutes of Health or American Diabetes Association.

## Author Contributions

M.L.S., M.E., F.S.C., D.U., S.C.J.P, Y.R., A.P.A., E.L.A., and P.S. designed the experimental and data analysis strategies. A.D. and E.P. performed spectral karyotyping. M.L.S., R.K., M.E., E.P., X.L., O.L., E.L.A., H.G., and Y.R., contributed to the generation and sequencing of EndoC-ßH1 and human islet libraries. N.L., A.T., E.J.M., P.O., S.C.J.P., A.V., M.K., P.S., M.E., P.S.C., S.R., M.S.S., and N.N. performed bioinformatics analyses of the data. Genotyping and imputation of EndoC-ßH1 was performed by M.E. and N.N. N.L., M.E., N.N., and M.L.S. wrote the manuscript and N.L., E.J.M., N.N., P.S., and M.K. designed the corresponding figures. All authors reviewed the manuscript prior to submission.

## Materials and Methods

### EndoC-ßH1 cell culture and processing

EndoC-ßH1 cells provided by EndoCells/INSERM were cultured and passaged as previously described (Ravassard et al., 2011). Briefly, cells were seeded at a density of approximately 600,000 cells/cm^2^ on tissue culture-treated plates pre-coated overnight with extracellular matrix (Sigma) and fibronectin (Sigma) in EndoC-ßH1 complete medium. Cells were passaged approximately every 7 days. Cells were harvested at various passages and distinct sites (e.g., NHGRI, JAX-GM) for karyotyping, genotyping, ATAC-seq, ChIP-seq, RNA-seq, Hi-C, and Pol2 ChIA-PET analyses.

### Human islet acquisition and procurement

The single human pancreatic islet sample used in this study was obtained from the National Disease Research Interchange (NDRI). The islet was shipped overnight from the distribution center. On receipt, we pre-warmed the islet to 37 °C in shipping media for 1–2 h before harvest; □50,000 islet equivalents (IEQs) were harvested for Hi-C.

### Spectral karyotyping (SKY)

Spectral karyotyping of EndoC-ßH1 was completed to identify structural and numerical chromosome aberrations using standard procedures as previously described. In brief, EndoC-ßH1 cells were cultured to 80% confluence. Metaphase spreads were prepared from these cells after mitotic arrest with Colcemid (0.015 μg/mL, 16 to 18 hours) (GIBCO, Gaithersburg, MD), hypotonic treatment (0.075 mol/L KCl, 20 minutes, 37°C), and fixation with methanol–acetic acid (3:1). Commercial SKY probe and software (Applied Spectral Imaging INC, Carlsbad, CA) was used to identify and visualize the individually colored chromosomes obtained from two slides’ worth of metaphase spreads from the same passage.

### Genotyping, imputation, and phasing of EndoC-ßH1

EndoC-ßH1 was genotyped with the HumanOmni2.5–4v1_H BeadChip Array (Illumina, San Diego, CA, USA). We mapped the Illumina array probe sequences to the hg19 genome assembly and excluded likely problematic ones as described in (Varshney et al., 2017).

We applied the following filtering criteria to remove additional SNP probes prior to pre-phasing of the array genotypes: 1) we assessed allele frequency of the SNPs using combined genotypes of EndoC-ßH1 and 163 other samples that were genotyped on similar chips; and 2) we removed SNPs with an alternate allele frequency difference with 1000G EUR samples > 20%, or palindromic SNPs with a minor allele frequency > 20%, genotype missingness > 2.5%, Hardy-Weinberg p-value < 10^−4^. At the end, a total of 1,851,388 SNPs were used in pre-phasing and imputation.

We performed pre-phasing and imputation separately on autosomal and chrX markers using the Michigan Imputation Server (Das et al., 2016). We used Eagle v2.3 (Loh et al., 2016) for autosomal chip marker pre-phasing and SHAPEIT v2.r790 (Delaneau et al., 2011) for chrX markers. We subsequently used minimac3 (Das et al., 2016) for imputation of missing genotypes using the Haplotype Reference Consortium (HRC version, hrc.r1.1.2016) panel (McCarthy et al., 2016).

### GWAS SNP pruning

Lists of reference SNP identifiers were obtained from the NHGRI-EBI Catalog of SNPs (**Error! Hyperlink reference not valid.**; accessed January 19^th^, 2017) for Type 2 diabetes, Type 1 diabetes, fasting glucose traits, fasting insulin traits, and proinsulin level categories. For each disease category, GWAS SNPs were pruned using PLINK version 1.9 (Purcell et al., 2007) to identify SNPs in high linkage disequilibrium (R^2^ > 0.8) as previously described (Lawlor et al., 2017b). T2D-associated SNPs from (Mahajan et al., 2018) were obtained and pruned using the same methodology described above.

### ATAC-seq

EndoC-ßH1 ATAC-seq libraries were prepared as previously described (Varshney et al., 2017) and sequenced on an Illumina NextSeq 500 with 2 × 125 bp cycles. Raw sequence fastq files for adipocyte tissue, bulk islet (Khetan et al., 2017), islet beta and alpha (GSE76268) (Ackermann et al., 2016), peripheral mononuclear blood cells (PBMC) (Ucar et al., 2017), skeletal muscle (Scott et al., 2016), GM12878 and CD4^+^ T cells (GSE47753) (Buenrostro et al., 2013) were obtained from their corresponding studies. Paired-end ATAC-seq reads were quality trimmed using *Trimmomatic* version 0.33 (Bolger et al., 2014) and parameters “TRAILING:3 SLIDINGWINDOW:4:15 MINLEN:36”. Trimmed reads were aligned to human genome (hg19) using BWA version 0.7.12 (Li, 2013), specifically using the bwa mem –M option. Duplicate reads were removed using “MarkDuplicates” from *Picard-tools* version 1.95 (The Broad Institute, 2013). After preprocessing and quality filtering, peaks were called on alignments with *MACS* version 2.1.0 (Zhang et al., 2008) using the parameters “-g ‘hs’ --nomodel --keep-dup all --broad --broad-cutoff 0.05 -f BAMPE”. Peaks located in blacklisted regions of the genome were removed. Remaining overlapping peaks from all cell types were merged with *BEDTools* version 2.26.0 (Quinlan and Hall, 2010) to generate a single peak set (n = 269,701). Raw read counts in these peaks for each cell type were determined using the R package *DiffBind_2.4.8* (Ross-Innes et al., 2012). Spearman rank-order correlation was calculated for cell types using the merged peaks with *deepTools* version 2.4.2 (Ramírez et al., 2014).

### ChIP-seq

CTCF, H3K27ac, H3K27me3, H3K36me3, H3K4me1, H3K79me2, H3K4me3, H3K9me3 ChIP-seq was performed as previously described (Stitzel et al., 2010) and sequenced on an Illumina HiSeq 2000 using 2 × 100 bp cycles. Harmonized ChromHMM (Ernst and Kellis, 2017) states for EndoC-ßH1 and NIH Roadmap cells/tissues were determined as previously described (Varshney et al., 2017).

### Transcription factor motif enrichment analysis

“findMotifsGenome.pl” (*HOMER* version 4.6 (Heinz et al., 2010)) script with parameters “hg19-size 200” was used to determine TF motifs enriched in ATAC-seq OCRs for each cell type (**Figure 2C**). In each analysis, all merged OCRs (n = 269,701) were provided as background (e.g., EndoC-ßH1 called OCRs (foreground) vs. all merged OCRs (background)). The same parameters were used to identify enriched motifs in either “Enhancers in Both EndoC-ßH1 and Islet” (n = 16,351) or “Enhancers in EndoC-ßH1 Only” (n = 19,380) compared to all enhancers (n = 51,325) (Figure 2D). The same HOMER script and parameters were also used to identify enriched motifs in EndoC-ßH1 (n = 315) vs. GM12878 (n = 308) cell-specific Hi-C loops.

### Similarity of cell/tissue type chromatin state (ChromHMM) annotations at EndoC-ßH1 ATAC-seq OCRs

EndoC-ßH1 ATAC-seq OCRs (n = 127,894) were overlapped with chromatin state (ChromHMM) annotations from EndoC-ßH1, human islet, adipocyte, skeletal muscle, GM12878, and PBMC cells provided in (Varshney et al., 2017). Next, only OCRs that intersected a ChromHMM annotation from all tissue/cell types (n = 127,887/127,894) were retained (union set). Within a tissue/cell type, or instances where multiple chromatin state elements intersected an EndoC-ßH1 OCR, annotations were prioritized as follows: promoter, enhancer, transcription, repressed, or low signal. At each OCR, cell/tissue ChromHMM annotations were compared to those of EndoC-ßH1 and assigned a binary classification (1 = the annotations were the same, 0 = the annotations were different). Aggregated counts of pairwise chromatin state annotations based on EndoC-ßH1 OCRs were then computed for each tissue/cell type, and a resulting similarity matrix was calculated using the “simil” function within the *proxy* version 0.4 R package (Meyer and Buchta, 2018).

### RNA-seq

Total RNA was extracted and purified from EndoC-ßH1 using Trizol as previously described (Varshney et al., 2017). All sequencing was performed on an Illumina NextSeq 500 with 2 × 101 bp cycles. Raw fastq files for human islets (Khetan et al., 2017), islet beta and alpha (Ackermann et al., 2016), PBMC (GSE90275), skeletal muscle (GSE78611), adipocyte (GSE93486) (ENCODE Project Consortium, 2012), GM12878 (GSE30400) (Rozowsky et al., 2011), and CD4^+^ T cell (GSE18927) (Schultz et al., 2015) were obtained from the associated databases. Paired-end RNA-seq reads were trimmed using *Trimmomatic* with the same parameters as used for ATAC-seq reads. Trimmed reads were aligned to human genome (hg19) using *STAR* version 2.53 (Dobin et al., 2012) with default parameters and expression levels of all genes were determined using *QoRTs* version 1.2.42 (Hartley and Mullikin, 2015) with default parameters and Gencode v19 transcript annotations. A total of 27,564 protein-coding genes and long intergenic non-coding RNAs (lincRNAs) were considered in the study.

### miRNA-seq

Total RNA was extracted and purified from 2000-3000 islet equivalents (IEQ) or 2 × 10^6^ EndoC-ßH1 cells using Trizol (Life Technologies). RNA quality was confirmed with Bioanalyzer 2100 (Agilent); islet samples with RNA integrity number (RIN) greater than 6.5 were prepared for miRNA sequencing; EndoC-ßH1 cells RNA RIN scores were > 9.0. miRNA libraries were prepared at the NIH Intramural Sequencing Core (NISC) from 1 µg total RNA using Illumina’s TruSeq Small RNA Library Kit according to the manufacturer’s guidelines, except a 10% acrylamide gel was used for better separation of library from adapters. Libraries were pooled in groups of about 8 for gel purification. Single-end 51 base sequencing was performed on Illumina HiSeq 2500 sequencers in Rapid Mode using version 2 chemistry. Data was processed using RTA version 1.18.64 and CASAVA 1.8.2. All resulting data was processed with miRquant 2.0 (Kanke et al., 2016).

### Hi-C

Hi-C libraries were generated as described in (Rao et al., 2014) and analyzed using the Juicer Tools version 1.75 pipeline (Durand et al., 2016a). We sequenced 6,065,763,792 Hi-C read pairs in EndoC-ßH1 cells, yielding 1,909,699,446 Hi-C contacts; we also sequenced 6,009,242,588 Hi-C read pairs in islet cells, yielding 1,516,995,339 Hi-C contacts. Loci were assigned to A and B compartments at 500 kB resolution. Loops were annotated using HiCCUPS at 5kB and 10kB resolutions with default Juicer parameters. This yielded a list of 9,100 loops in EndoC-ßH1 cells and 2,580 loops in Islet cells. GM12878 loop calls (n = 9,448 loops) were downloaded from Gene Expression Omnibus (GSE63525). Differential loop calling with HiCCUPS at 5kb and 10kb identified 1,120 loops as significantly enriched for EndoC-ßH1 cells and 829 loops as significantly enriched for GM12878 cells. Similar comparison of islet and EndoC-ßH1 loops determined 935 loops as significantly enriched for EndoC-ßH1 and 49 loops as being significantly enriched for islet. Aggregate peak analysis (APA) plots were calculated using Juicer and the “apa” command using default parameters. Visualization of Hi-C maps was performed using Juicebox version 1.6.11 (Durand et al., 2016b) with the “Observed/Expected” view and “Balanced” (Knight-Ruiz) normalization. All the code used in the above steps is publicly available at (github.com/theaidenlab). Genomic Regions Enrichment of Annotations Tool (GREAT; (McLean et al., 2010) was used to identify pathways enriched in the single nearest genes (whose TSS was within 2 kb) of EndoC-ßH1-specific anchors.

### ChIA-PET

EndoC-ßH1 RNA Polymerase 2 (Pol2) ChIA-PET libraries were generated and sequenced reads were processed and analyzed according to the protocol in (Li et al., 2017b). ChIA-PET interactions were identified using ChIA-PET2 (Li et al., 2017a) using the “bridge linker mode” option. Corresponding ChIA-PET interactions for K562 (GSE39495) and GM12878 (GSE72816) cells were obtained from Gene Expression Omnibus. ChIA-PET and Hi-C loops were further filtered using the Bioconductor package *InteractionSet_1.8.0* (Lun et al., 2016) to retain only those in which both interacting sites (anchors) overlapped OCRs. ChIA-PET anchors were annotated to the nearest gene. To assess the physical connectivity between genes and their putative regulatory regions as captured by ChIA-PET interactions, the number of distinct links between anchors annotated to each gene were counted and categorized by their chromatin state and regulatory function. Counting was carried out both between each gene promoter and all linked regulatory regions (Fig. 4C), and also between all annotated anchors regardless of their chromatin state and all linked regulatory regions (Fig. S4B). For example, consider anchors A1 and A2, both overlapping enhancer regions annotated to gene *i*. These anchors respectively link to anchors B1, located in a TSS, and B2, in an enhancer. In this scenario, a connectivity degree of two would be computed for gene *i*, corresponding to an enhancer-TSS and a TSS-TSS interaction, respectively.

The functional specificity of EndoC-ßH1 ChIA-PET interactions was investigated by overlapping interaction anchors on ChromHMM chromatin states computed from EndoC-ßH1 data as well as 27 other tissue/cell types (Varshney et al., 2017), and calculating the rate of conservation of chromatin states of both anchors. For example, a rate of 80% enhancer-enhancer conservation would mean that 8 out of 10 interactions of this type in EndoC-ßH1 are also found in another cell type. The resulting proportions were computed for all interactions combined, and for specific relevant regulatory interactions (Figure 4D, line plots). In addition, aggregated counts of pairwise chromatin state interactions based on EndoC-ßH1 ChIA-PET interactions were computed for the same cell types as above, and pairwise distances (*D*) between the resulting 29 count matrices were computed and plotted as scaled similarity values relative to EndoC-ßH1 (i.e., 1-*D*/*D*_max_), so that *D* = 0 for EndoC-ßH1 interactions and *D* = D_max_ for the most divergent cell type (Figure 4D, bar plot). These same methods were used to determine the functional specificity of EndoC-ßH1 Hi-C interactions.

### Visualization of multiomic data

Multiomic plots of all Hi-C, ChIA-PET, chromatin state (ChromHMM), ATAC-seq, and RNA-seq data examined in this study were produced using the Bioconductor package *Sushi_1.18.0* (Phanstiel et al., 2014).

### ATAC-seq allelic bias analysis

All allelic bias analyses were performed using WASP (Geijn et al., 2015) (version 0.2.2 after GitHub commit 5a52185 and bug fix in pull request #67). For the EndoC-ßH1 ATAC-seq allelic bias analyses, after original BWA mapping, reads were filtered to properly-paired, high-quality autosomal reads using SAMtools (v. 1.3.1; flags -f 3 -F 4 -F 8 -F 256 -F 2048 -q 30) (Li et al., 2009). Remapping and filtering as part of the WASP pipeline utilized the same parameters. As the last step of the WASP pipeline, duplicate removal was performed using WASP’s rmdup_pe.py script. In order to avoid double-counting SNPs covered by both reads in a pair, overlapping read pairs were clipped using bamUtil’s clipOverlap (https://genome.sph.umich.edu/wiki/BamUtil:_clipOverlap; v. 1.0.14). The four replicate libraries were then merged using SAMtools merge.

For each SNP, we determined the number of reads containing each allele (requiring base quality of at least 20). We excluded SNPs with total coverage less than 10, as well as SNPs in regions blacklisted by the ENCODE Consortium because of poor mappability (wgEncodeDacMapabilityConsensusExcludable.bed and wgEncodeDukeMapabilityRegionsExcludable.bed). Allelic bias testing was performed using a two-tailed binomial test, using an adjusted expectation for the null to account for residual reference bias as described in (Scott et al., 2016). Briefly, for each of the 16 reference-alternate allele pairs (e.g., AG and GA are separate allele pairs), we calculated the expected fraction of reference alleles (fracRef) under the null as the sum of the reference allele counts divided by the sum of the total allele counts for SNPs of that allele pair. To prevent SNPs of high coverage from biasing the expected fracRef, we down-sampled SNPs with coverage in the top 25th percentile to the median coverage, and used these downsampled reference and total allele counts when calculating the expected fracRef. We used the observed allele-pair specific fracRef as the true fracRef under the null hypothesis of no allelic bias in the binomial test. Multiple testing correction was performed using the Benjamini-Hochberg correction (FDR < 10%).

### ChIP-seq allelic bias analysis

For the EndoC-ßH1 ChIP-seq allelic bias analyses, paired-end libraries were processed as follows. Adapters were trimmed using cta (v. 0.1.2) and reads mapped using BWA mem (-M flag; v. 0.7.12). Reads were filtered to properly-paired, high-quality autosomal reads using SAMtools (flags -f 3 -F 4 -F 8 -F 256 -F 2048 -q 30). Single-end libraries were mapped using BWA aln (v. 0.7.12; default parameters) and filtered using SAMtools (flags -F 4 -F 256 -F 2048 -q 30). Remapping and filtering as part of the WASP pipeline utilized the same parameters as for the original mapping. For paired-end libraries, overlapping read pairs were clipped using bamUtil’s clipOverlap. Replicates were merged using SAMtools merge. Allele counting and allelic bias testing was performed as described above for ATAC-seq.

### RNA-seq allelic bias analysis

For the EndoC-ßH1 RNA-seq allelic bias analyses, after original STAR mapping, reads were filtered to properly-paired, high-quality autosomal reads using SAMtools (v. 1.3.1; flags -f 3 -F 4 -F 8 -F 256 -F 2048 -q 255). Remapping and filtering as part of the WASP pipeline utilized the same parameters. In order to avoid double-counting SNPs covered by both reads in a pair, overlapping read pairs were clipped using bamUtil’s clipOverlap. Allele counting and allelic bias testing was performed as described above for ATAC-seq.

### Comparison of islet eQTL and EndoC-ßH1 biased SNPs allelic effect

Human islet eQTL data were obtained from (Varshney et al., 2017). For EndoC-ßH1 biased (H3K27ac) enhancer SNPs that were also linked (via ChIA-PET chromatin interaction) to a target gene (n = 91/185 SNPs in Figure 5C), all corresponding islet eQTL SNP-gene pairs were retrieved. For 42/91 SNPs linked to target genes, the H3K27ac allelic effect bias was calculated assuming that the EndoC-ßH1 effect allele was the same as the islet eQTL effect allele. Allelic effect bias was calculated by dividing the effect allele coverage (either reference or alternate allele) by the total coverage of the SNP. A scalar value of 0.5 was subtracted from this value to determine whether the effect allele had an increased (positive value), decreased (negative value), or no (zero) bias in H3K27ac coverage. Randomly selected eQTL SNP-gene pairs that did not have corresponding connections (via ChIA-PET chromatin interaction) to a target gene were considered as a null/background set.

## EndoC-ßH1 Supplementary figure legends

**Figure S1:**

Representative spectral karyotypes (SKY) of EndoC-ßH1 cells at metaphase. Specific metaphases shown are S2.1 (top), 7 (middle), and S2.7 (bottom). Common structural or numerical chromosomal aberrations that are evident in multiple metaphases are indicated in green.

**Figure S2:**

(A) Similarity (centered Pearson correlation) of human islet, adipocyte, skeletal muscle, GM12878, and PBMC chromatin state annotations (ChromHMM) to those of EndoC-ßH1 at EndoC-ßH1 OCRs. The similarity matrix was calculated using the “simil” function within the *proxy* version 0.4 R package (Meyer and Buchta, 2018). (B) Spearman correlation between EndoC-ßH1 RNA-seq and the corresponding RNA-seq from islets, sorted alpha or beta cells (Ackermann et al., 2016), and other cell types and tissues. PBMC=peripheral blood mononuclear cells, GM12878 = lymphoblastoid cell line, CD4T = CD4^+^ T immune cell, skeletal = skeletal muscle, Alpha = primary islet alpha cells, Beta = primary islet beta cells. (C) Principal component analysis (PCA) of miRNA-seq profiles from EndoC-ßH1 and 5 representative human islet, skeletal muscle, and adipose tissue samples. (D) Scatter plot illustrating the resemblance of miRNA expression levels between EndoC-ßH1 and human islets. RPMMM = reads per million mapped miRNA, R^2^ = Pearson R.

**Figure S3:**

(A) Biological process gene ontology terms enriched in genes adjacent to EndoC-ßH1-specific Hi-C anchors. Enrichment analysis was performed with GREAT (McLean et al., 2010) using the single gene whose transcription start site (TSS) was nearest to each anchor. Results with an adjusted (Benjamini & Hochberg) hypergeometric p-value < 0.05 were regarded as statistically significant. (B) Frequency of EndoC-ßH1 Hi-C loop anchors and their corresponding chromatin state (ChromHMM) annotations. (C) (Top) Bar plot depicting the average similarity of chromatin state (ChromHMM) annotations for each cell type at EndoC-ßH1 Hi-C loop anchor positions. Overlaid line plots highlight the relative similarity for various chromatin state interactions (e.g. active promoter vs. active promoter). (Bottom) Select heat maps showing the frequency of human islet (left) and GM12878 (right) chromatin states at EndoC-ßH1 loop anchors.

**Figure S4:**

(A) Genome-wide view of RNA polymerase 2-mediated (Pol2 ChIA-PET) chromatin interactions around the *PDX1* locus on chromosome 13 in EndoC-ßH1. Asterisks under EndoC-ßH1 (red) ChIA-PET interactions indicate interacting sites/anchors identified by targeted 4C-seq analyses in this locus in human islets (Pasquali et al., 2014). (B) Connectivity of EndoC-ßH1 ChIA-PET interactions when considering nodes with 6 or more links to other regulatory elements. (Left) Circular plots depict the number of links that occur with corresponding regulatory elements (e.g., active promoter, active enhancer). (Right) bar plots illustrate the proportion of interacting nodes that exhibit the active regulatory element chromatin states exclusively in EndoC-ßH1 (blue) or identical chromatin states in both EndoC-ßH1 and islet (green). (C) Multidimensional scaling (MDS) plot based on pairwise Chi-square distances of vectors of proportions of chromatin states in EndoC-ßH1, islets, and additional Epigenomics Roadmap cell and tissue types at EndoC-ßH1 defined ChIA-PET interacting nodes.

**Figure S5:**

(A) Correlation of *cis*-regulatory element allelic imbalance for SNPs with heterozygous genotypes in EndoC-ßH1. For a given heterozygous SNP (n = 1,734 total with 20X coverage in EndoC-ßH1 ATAC-seq and H327ac data), allelic imbalance ratios were calculated and plotted. Colored points signify SNPs with significant allelic imbalance (FDR < 10%) in ATAC-seq (red), H3K27ac (blue), or both (purple) datasets. (B) Correlation of human islet eQTL SNP-gene pair direction of effect (z-score) from (Varshney et al., 2017) and their corresponding H3K27ac effect allele bias deviation from 0.5 in EndoC-ßH1. Red points indicate eQTL SNP-gene pairs that are also linked by an EndoC-ßH1 ChIA-PET interaction. Asterisks indicate eQTL SNPs that are also diabetes-associated GWAS SNPs.

